# Voltage imaging using transgenic mouse lines expressing the GEVI ArcLight in two olfactory cell types

**DOI:** 10.1101/2020.08.26.268904

**Authors:** Jelena Platisa, Hongkui Zeng, Linda Madisen, Lawrence B. Cohen, Vincent A Pieribone, Douglas A. Storace

## Abstract

Genetically encoded voltage indicators (GEVIs) allow for cell-specific optical recordings of membrane potential changes in defined cell populations. One tool that would further their use in the *in vivo* mammalian brain is transgenic reporter animals that facilitate precise and repeatable targeting with high expression levels. The present literature on the development and use of transgenic mouse lines as vehicles for GEVI expression is limited. Here we report the first *in vivo* experiments using a transgenic reporter mouse for the GEVI ArcLight (Ai86(TITL-ArcLight)), which utilizes a Cre/tTA dependent expression system (TIGRE 1.0). Following pairing to appropriate Cre- and tTA transgenic mice, we report two mouse lines with ArcLight expression restricted to olfactory sensory neurons (OMP-ArcLight), and a subpopulation of interneurons that include periglomerular and granule cells (Emx1-ArcLight) in the olfactory bulb (OB). The ArcLight expression in these lines was sufficient for *in vivo* imaging of odorant responses in single trials. Odor responses were measured in the OB using epifluorescence and 2-photon imaging. The voltage responses were odor-specific and concentration-dependent, and confirmed earlier conclusions from calcium measurements. This study shows that the ArcLight Ai86(TITL-ArcLight) transgenic line is a flexible genetic tool that can be used to record neuronal electrical activity of a variety of cell types with a signal-to-noise ratio that is comparable to previous reports using viral transduction.

## Introduction

Genetically encoded voltage indicators (GEVIs) can report changes in membrane potential from genetically defined cell-types. The membrane localization, and the spatial and temporal complexity of these signals make the design and production of GEVIs challenging [1–6]. However, recent advances in GEVI development have resulted in improvements in the sensitivity, speed and signal-to-noise ratio in several GEVIs, allowing for enhanced detection of electrical transients *in vivo* [7–17].

While sensitivity is critically important, widespread adoption of GEVIs for *in vivo* experiments would benefit from the availability of tools that facilitate selective expression in specific cell types. To date, most mammalian *in vivo* approaches have driven expression using *in utero* electroporation [12, 17] or injection of a viral vector (e.g. adeno-associated viruses, AAVs) [10, 11]. These protocols complicate the experiment by requiring additional surgical procedures, and in some instances, available viruses have limited transduction capacity for particular cell types (e.g., olfactory receptor neurons). Transgenic animals can overcome these limitations as demonstrated by studies using GEVIs in *Drosophila* where the use of bipartite systems such as GAL4/UAS allow for targeted expression in any brain region [18–20].

However, there are currently a limited number of transgenic mice expressing GEVIs with high signal-to-noise ratios (Butterfly 2.1 [21], QuasAr2 [22], ASAP2s [23]), and even fewer studies showing their utility *in vivo* [24, 25]. Here we took advantage of a novel transgenic reporter mouse, Ai86(TITL-ArcLight) [21, 23], that can be used to selectively drive expression of the GEVI ArcLight [7] in specific cell types. This transgenic line expresses ArcLight in the presence of both tetracycline transactivator (tTA) and Cre recombinase [21, 26].

The olfactory bulb (OB) is the first region of information processing in the mouse olfactory system. The olfactory sensory neuron (input) projections, and the dendrites of the mitral/tufted projection neurons (output) both innervate OB glomeruli. GEVIs have been used to study sensory processing in different OB cell types using viral transduction as the expression vector [10, 11]. However, results from our laboratory have shown that the tested AAVs had limited transduction efficacy for olfactory sensory neurons (D. A. S. and L. B. C, unpublished observations). Here we report the generation of a novel transgenic line with ArcLight expression localized to these sensory neurons (Ai86 (TITL-ArcLight) x CamkII-tTA x OMP-Cre) called OMP-ArcLight, and demonstrate its ability to report odor-evoked signals *in vivo*. We also report a second transgenic line that expresses ArcLight primarily in OB interneurons (Ai86 (TITL ArcLight) x CamkII-tTA x Emx1-Cre) called Emx1-ArcLight.

## Material and methods

### Transgenic mice

All described experiments were performed in strict accordance with the recommendations in the Guide for the Care and Use of Laboratory Animals of the National Institutes of Health. The protocol used was approved by the Institutional Animal Care and Use Committees at Yale University and The John B Pierce Laboratory.

The ArcLight transgenic mouse line, Ai86(TITL-ArcLight) (Ai86; JAX Stock No. 034694), was generated by the Allen Institute for Brain Science [21, 23]. The detailed description of the mouse line and the intersectional transgenic approach can be found in [21]. Ai86 is a Cre/tTA-double reporter mouse line in which the GEVI ArcLight A242 [7] is knocked into the TIGRE locus (TIGRE 1.0). For this study, the Ai86(TITL-ArcLight) founder line was independently crossed to either Camk2a-tTa (Jax Stock #007004) or Emx1-Cre (Jax Stock # 005628) yielding double mutant lines Ai86 x CamK2a-tTA and Ai86 x Emx1-Cre, respectively. The transgenic offsprings of these lines were crossed to create a triple transgenic mouse line Ai86 x Camk2a-tTa x EMX1-Cre (named Emx1-ArcLight) with ArcLight expression targeted to the granule cell layer and a small population of periglomerula cells in the OB. Out of 178 pups produced in these crossings 28 were triple transgenic (15.7%). An additional triple transgenic line with ArcLight targeted to the olfactory sensory neurons was crated by crossing Ai86 x Camk2a-tTA and OMP-Cre (Jax Stock # 006668), lines yielding Ai86 x Camk2a-tTa x OMP-Cre (named OMP-ArcLight). Out of 45 pups produced in these crossings 11 were triple transgenic (24.4%). The conf**i**rmation of presence of ArcLight, tTA, and Cre DNA was done via PCR-based genotyping performed either in house or by Transnetyx (Cordova, USA) on all experimental animals. Mice were housed under standard environmental conditions, with room temperature ranging between 23-25°C and under a 12h light/dark cycle. Our measurements were made during the light phase. Adult mice were examined for crude physical or behavioral abnormalities.

### Histology and confocal imaging

Following imaging, mice were euthanized (Euthasol®) and brains (Emx1-ArcLight: N = 5; OMP-ArcLight: N = 3) were dissected and fixed in 4% paraformaldehyde for a minimum of 3 days. The OBs were embedded in 3% agarose, and cut on a vibratome in 50–70 μm thick coronal slices. Sections were mounted on microscope slides using VECTASHIELD Mounting Medium with DAPI (Vector Labs, H-1500). Confocal images were obtained using a Zeiss LSM-780 confocal microscope (Carl Zeiss Microsystems, USA). Fluorescence was visually evident without any amplification procedures, and thus all histological fluorescence is the result of endogenous expression.

### Surgical procedure

All surgical procedures were performed under sterile conditions. Male and female mice agese between 30-180 days were used for functional imaging experiments. Ketamine/xylazine (90 mg kg^−1^/10 mg kg^−1^) (Covetrus, USA) were used to induce and maintain a deep anesthetic state via intraperitoneal (IP) injections. Anesthetic depth was frequently assessed via the pedal reflex. A heating pad was used for maintenance of body temperature at 37°C throughout the procedure. Lidocaine (0.5%) was injected at the wound area. IP injections of atropine (Covertus, USA) (0.2 mg/kg) were administered or prevention of excessive bronchial secretion. The head was shaved and scrubbed with Betadine. The skin above the cranium and OB was removed, and a custom head-post was fixed to the back of the skull using either cyanoacrylate or Metabond (C&M Metabond, Parkell, USA). The mouse was then mounted on a custom-made holder, which allowed for precise head positioning and fixation. Depending on the experiment, a high-speed dental drill (XL-230 Osada, Japan) was used to either thin or remove the skull above both OBs. For 2-photon imaging, the craniotomy above the bulb’s surface was covered with 2% agarose and sealed with a #1 glass coverslip.

### Odorant stimuli and delivery

Different odorants (methyl valerate, isoamyl acetate, ethyl tiglate, and 2-heptanone; Sigma-Aldrich, USA) were used at concentrations between 0.12% and 11% of saturated vapor. A cleaned air stream was used for odorant dilution from saturated vapor. A flow dilution olfactometer [27] was designed to provide a constant airflow over the nares. Odorant delivery was controlled by a vacuum suction, with the vacuum switched off during the odorant presentation. Separate teflon tubing lines were used for each odor to avoid cross-contamination. The time course and relative concentration of odors were monitored with a photo-ionization detector (PID; Aurora Scientific, Aurora, ON). In a subset of experiments, the PID placed next to the mouse’s nose in front of the olfactometer to confirm the odor timecourse. In some imaging trials odorants were delivered for 2-3 seconds, with a 60 second delay. In other trials, the odor was repeatedly presented with a 6-s interstimulus-interval.

### Imaging systems

The epifluorescence imaging was performed on a custom upright microscope equipped with a Prizmatix LED (UHP-T-LED-White-High-CRI), using either of two objectives a 35 mm F/1.4 Computar or a 25 mm F/0.95 Computar CCTV lens. We used 479 nm (Semrock FF01-479/40) excitation filter combined with a 515 nm long pass dichroic mirror, and a bandpass filter. The neuronal activity was recorded with a NeuroCCD-SM256 camera (RedShirtImaging, USA) with 2 × 2 binning and with a frame rate between 50–250 Hz. The images were collected with NeuroPlex software (RedShirtImaging, USA).

For 2-photon imaging, we used a modified MOM two-photon laser-scanning microscope (Sutter Instruments, USA), a Coherent Discovery laser light source, and detected fluorescence emission on GaAsP PMTs (#H10770PA-40-04, Hamamatsu, Japan). The excitation of the super ecliptic pHluorin GFP chromophore was achieved with 940-980 nm laser light with imaging speed of 31 frames per second (resonant scanners; Cambridge Technology, USA). The laser power at the surface of the sample was between 75 to140 mW.

### Data analysis

The data analysis was done in NeuroPlex and Excel. The presented optical traces are the spatial average of all the pixels within a region of interest (ROI). For the OMP-ArcLight line, individual glomeruli were visually identified as circular peaks of activation ~50–100 μm in diameter. Response amplitudes for identified glomeruli were identified as the difference in the temporal average of the 1–2 s preceding the stimulus from a 0.8–1 s average around the peak of the response. Data are presented as the change in fluorescence divided by the resting fluorescence, ΔF/F. When necessary, the fluorescence traces were corrected to remove photobleaching by dividing the signal by a single exponential curve fitted to the portion of the trace prior to the stimulus. ArcLight reports voltage depolarization as decreases in fluorescence [7], and traces are inverted such that depolarizations are shown upwards. Pixels contaminated with blood vessel artifacts in major vessels were omitted from the analysis.

The activation frame subtraction maps were generated in NeuroPlex (Frame Subtraction feature) by subtracting the temporal average of the 1–2 s preceding the stimulus from a 1 s temporal average around the response peak. No spatial filtering was performed; the activity maps are displayed as depixelated (Depixelation function in NeuroPlex).

The error bars in Figures 2 and 4 and the statistics in the Results represent the standard error of the mean (s.e.m.).

## Results

*In vivo* imaging was carried out in two transgenic lines in which ArcLight was targeted to two different cell populations using the transgenic strategy shown in Figure 1A. These reporter mice utilize an intersectional tTA/Cre TIGRE 1.0 system in which ArcLight is selectively expressed in cells that also express both tTA and Cre [21, 23]. A diagram of the breeding schemes used to generate the two lines is shown in Figure 1B. The Ai86 (TITL-ArcLight) line was crossed to either the CamkIIa-tTA or Emx1-Cre mouse line, resulting in two new lines that either expressed ArcLight and tTA (ArcLight-tTA), or ArcLight and Emx1-Cre (ArcLight-Emx1) (Figure 1B). The cross between these lines resulted in offspring with ArcLight expression in a population of olfactory bulb interneurons (named Emx1-ArcLight). An additional line (named OMP-ArcLight) that expressed ArcLight in the olfactory sensory neurons was generated by crossing the ArcLight-tTA and OMP-Cre lines.

**Figure 1.**
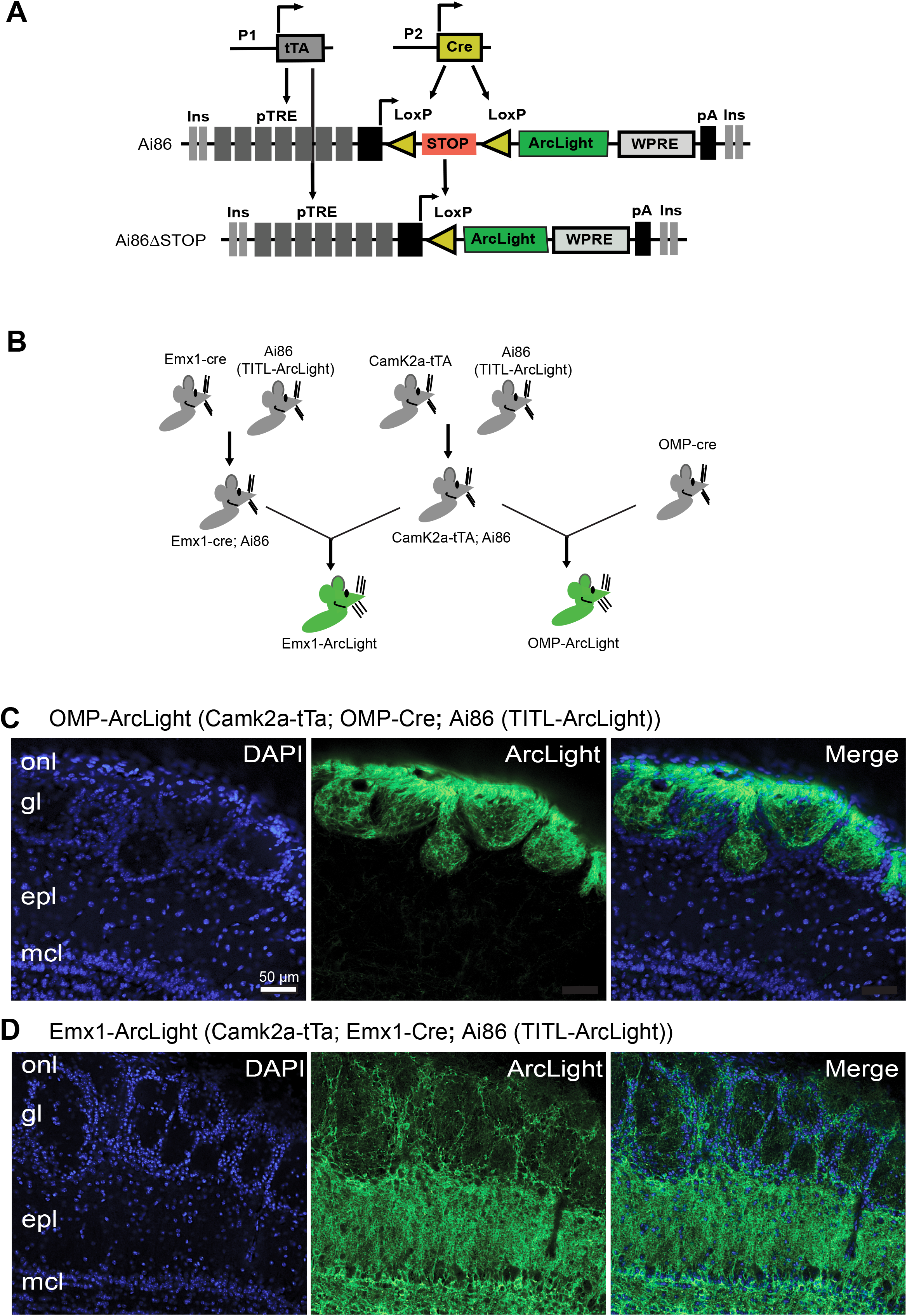
Expression of ArcLight in the olfactory bulb of transgenic mouse lines is robust and cell-specific. **A.** The design of the reporter line, Ai86 (TITL-ArcLight), is based on the Cre and tTA-dependent intersectional TIGRE1.0 approach. **B**. Breeding scheme used to create two triple transgenic mouse lines that show cell-specific expression of GEVI ArcLight in the olfactory bulb. **C.** Coronal slices of olfactory bulb were imaged using confocal microscopy. The images show membrane-localized expression of GEVI ArcLight targeted to olfactory receptor neurons in the OMP-ArcLight mouse. **D.** In the EMX-ArcLight transgenic mice, ArcLight is mainly targeted to the olfactory bulb's glomerular cell layer. In both C. and D., DAPI (left), ArcLight (middle), and merge (right). In all images, the scale bar size is 50 μm. onl, olfactory nerve layer; gl, glomerular layer; epl, external plexiform layer; mcl, mitral cell layer.

### Robust and cell-specific expression of ArcLight in transgenic mouse lines

We confirmed the expression pattern of ArcLight in both transgenic lines in histological sections. The OMP-ArcLight line (3 preparations) had fluorescence restricted exclusively to the olfactory nerve and glomerular layer (e.g., Figure 1C). The Emx1-ArcLight line (5 preparations) had fluorescence localized to the granule cell layer, and sparse labeling of individual neurons in the glomerular layer (e.g., Figure 1D) [28]. As previously reported [23], the cortical pan-expression of ArcLight in these mice was confirmed (data not shown). The odorant responses in both mouse lines were detectible in a single trial, and the signal was not diminishing during prolonged imaging sessions (up to 2 hours).

### OMP-ArcLight mice

#### ArcLight can detect odor dependent activity in a single trial

Responses to different odor-concentrations were imaged across the dorsal OB using epifluorescence imaging. We detected odors evoked glomerular sized peaks of activity in a frame subtraction analysis that were odor- and concentration dependent (between 3-11 glomeruli per preparation, N = 7 preparations; Figure 2A). Different odors evoked distinct patterns (Figure 2A) are in agreement with prior reports that imaged olfactory sensory neurons (OSNs) using sensors of calcium or synaptic vesicle release [29, 30].

**Figure 2.**
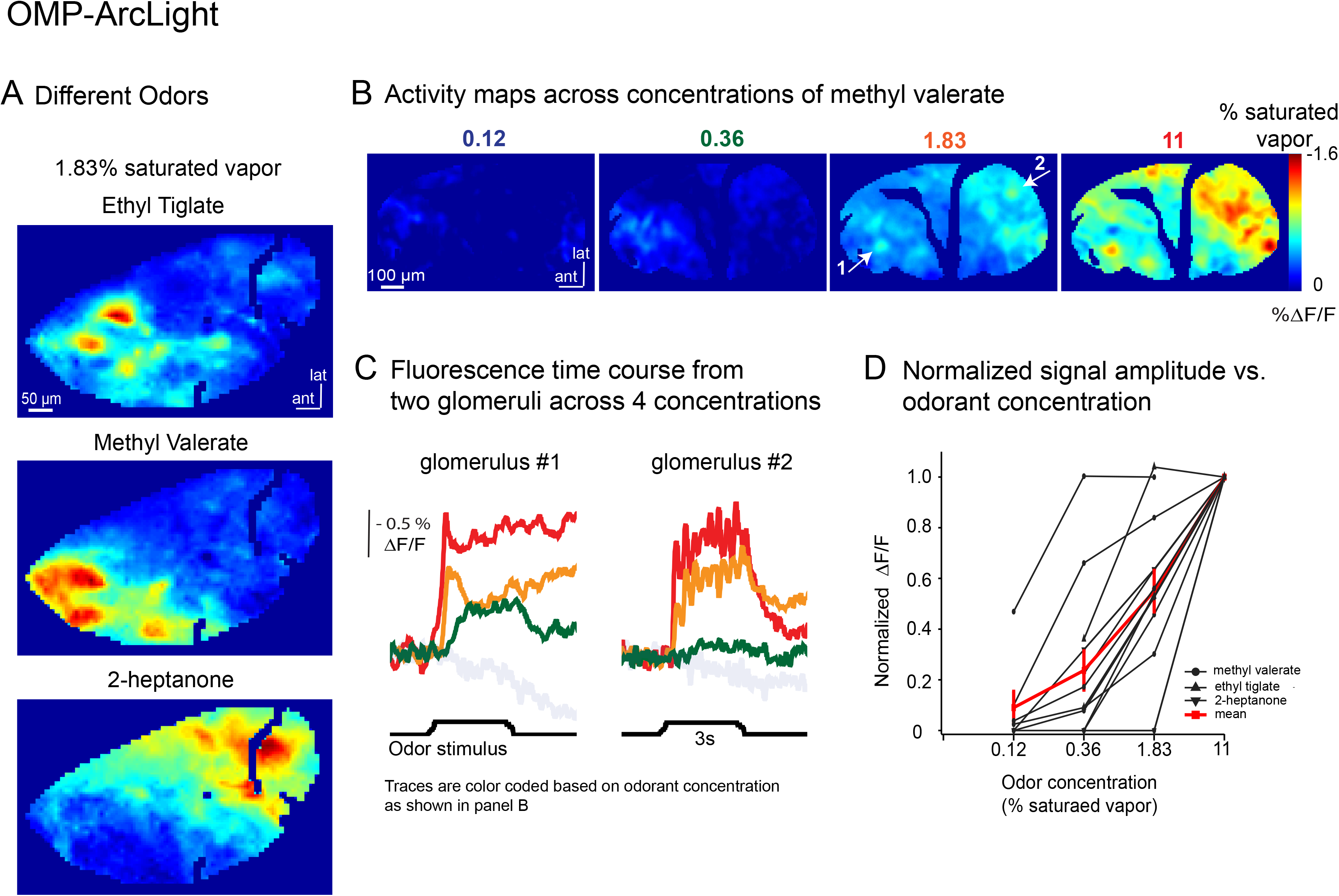
Odor-evoked activity in the olfactory receptor neurons (input) of the OMP-ArcLight mice recorded with the GEVI ArcLight. **A**. Activity maps show odorant-specific glomerular activity. Three odorants were tested: ethyl taligate (upper), methyl valerate (middle), and 2-heptanone (lower). All three odorants were administrated at the concentration of 1.83% saturated vapor. For each odorant, the minimum and maximum %ΔF/F values are shown in the panel. Each activity map is an average of three recordings aligned to the first respiration. Bar size 50μm. **B**. A series of activity maps show the concentration-dependent response in the olfactory bulb’s receptor neurons activity. The four concentrations of methyl valerate used are indicated in color-coded numbers above each panel. The color scale on the right shows the range of %ΔF/F values from minimum to maximum. The bar size 100 μm. **C.** Examples of optical traces of odor-evoked activity in response to a single stimulus (3s long) at three different concentrations of the odorant methyl valerate. The responses were measured from two glomeruli shown in panel B (white arrows in the 1.83% panel). The single trial, non-low pass filtered traces are color-coded depending on odor concentration as in panel B. **D**. The normalized signal amplitude vs. odorant concentration shows a sharp dependence of neuronal response on odor concentration. The responses for the lower odorant concentrations were normalized to responses recorded at the highest concentration (11% saturated vapor). The responses were recorded from 71 glomeruli in seven animals to three different odorants (grey lines). The average of all responses is shown as a red line, with s.e.m as error bars. The glomeruli were defined based on the response to 1.83% of saturated vapor.

#### Concentration dependence

Increasing the odor concentration caused increases in both the number of activated glomeruli (Figure 2B) and overall signal size (Figure 2C, D). Exemplar single trial traces from two glomeruli in response to three different odor concentrations are shown in Figure 2C. The concentration-response function for a population of preparations is plotted in Figure 2D. The responses to lower odorant concentrations (0.12 %, 0.36 %, and 1.83 % of saturated vapor) were normalized to the highest concentration response (11 % of saturated vapor). The average signal size of all the glomeruli activated for each preparation is plotted in Figure 2D (thin black lines), along with the average of all the preparations (thick red line +/− s.e.m. The concentration-dependent increase in the response amplitude for the olfactory receptor neuron input was relatively steep, consistent with the prior reports from calcium measurements [11, 30, 31]. For 2-heptanone and ethyl tiglate, the lowest concentration of odorant used was 0.36 % of saturated vapor. The response to the lowest concentration was ~8.36% of the response to the highest concentration.

#### Different temporal patterns of activity across the olfactory bulb glomeruli

Odor stimulation evoked activity in glomeruli with distinct temporal characteristics [32]. The activity map evoked by an odor (methyl valerate) across the 2 s odor-presentation is shown in Figure 3A. Single trial recordings from a caudal and rostral glomerulus demonstrate dramatically different temporal properties (Figure 3B), consistent with prior reports [10, 32]. The signal from the caudal glomerulus returns part of the way to the resting level between inhalations and is modulated by respiration. In contrast, the rostral glomerulus was slower to respond and had less respiratory modulation (Figure 3B).

**Figure 3.**
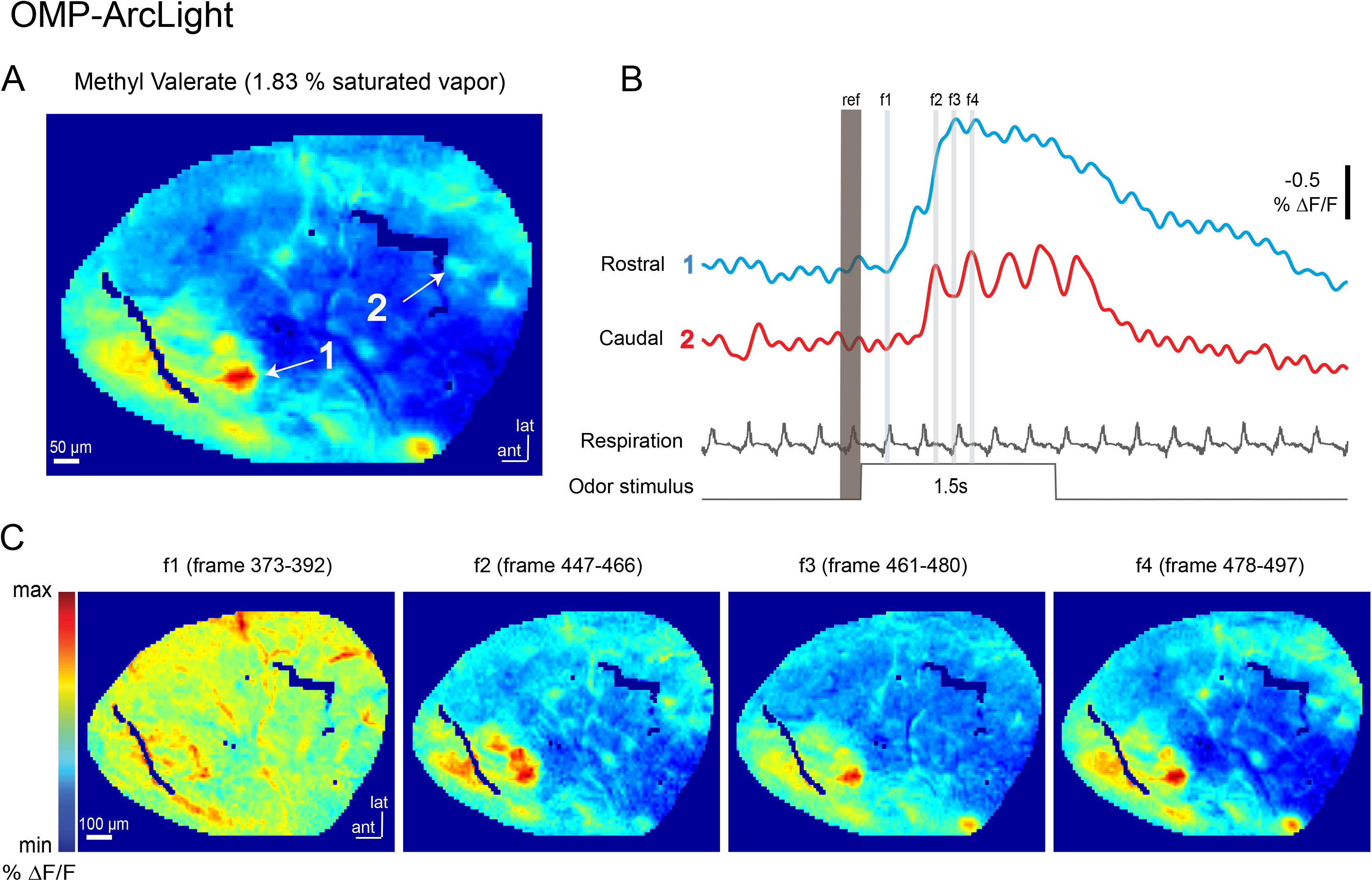
Temporally diverse responses detected with the GEVI ArcLight in the olfactory bulb input of the OMP-ArcLight transgenic mouse line. **A.** Activity map of neuronal activity in response to 1.83% saturated vapor of the odorant (methyl valerate) with examplar glomeruli located in the rostral and caudal part of the olfactory bulb (white arrows). **B.** Optical traces filtered with a 6Hz low-pass filter show diverse odor-evoked activity of ArcLight detected in the two different glomeruli designated in panel A. **C.** Activity maps created at the different time points (f1-f4 in panel B) during odor stimulus show differences in the temporal dynamics of response recorded in the two glomeruli shown in panel A and B. The color scale on the right shows the range of %ΔF/F values from minimum to maximum. Bar size 100 μm.

These respiratory driven dynamics are visualized via activity maps evoked at different time points during the odor period (Figure 3C, f1-f4).

#### Adaptation in the olfactory bulb input

We examined the stability of OSN odor responses by measuring the response to repeated odor presentations of 3-s separated by a 6-s interstimulus interval. An example of an activity map with two glomeruli (white arrows) activated by 11% saturated vapor of methyl valerate is shown in Figure 4A. Single trial optical responses from two glomeruli from trials in response to 1.83% and 11% of saturated vapor are shown in Figure 4B. The measurements showed minimal adaptation across repeated stimuli in both glomeruli. The analysis across all preparations and odorants confirmed this trend (Figure 4C). The response amplitude to the 2nd and 3rd odor presentation was normalized to the initial response amplitude (Figure 4C). The average glomerular response for each preparation is shown as the thin black trace, and the population mean in red (+/− s.e.m.; 3-11 glomeruli per prep; N = 7).

**Figure 4.**
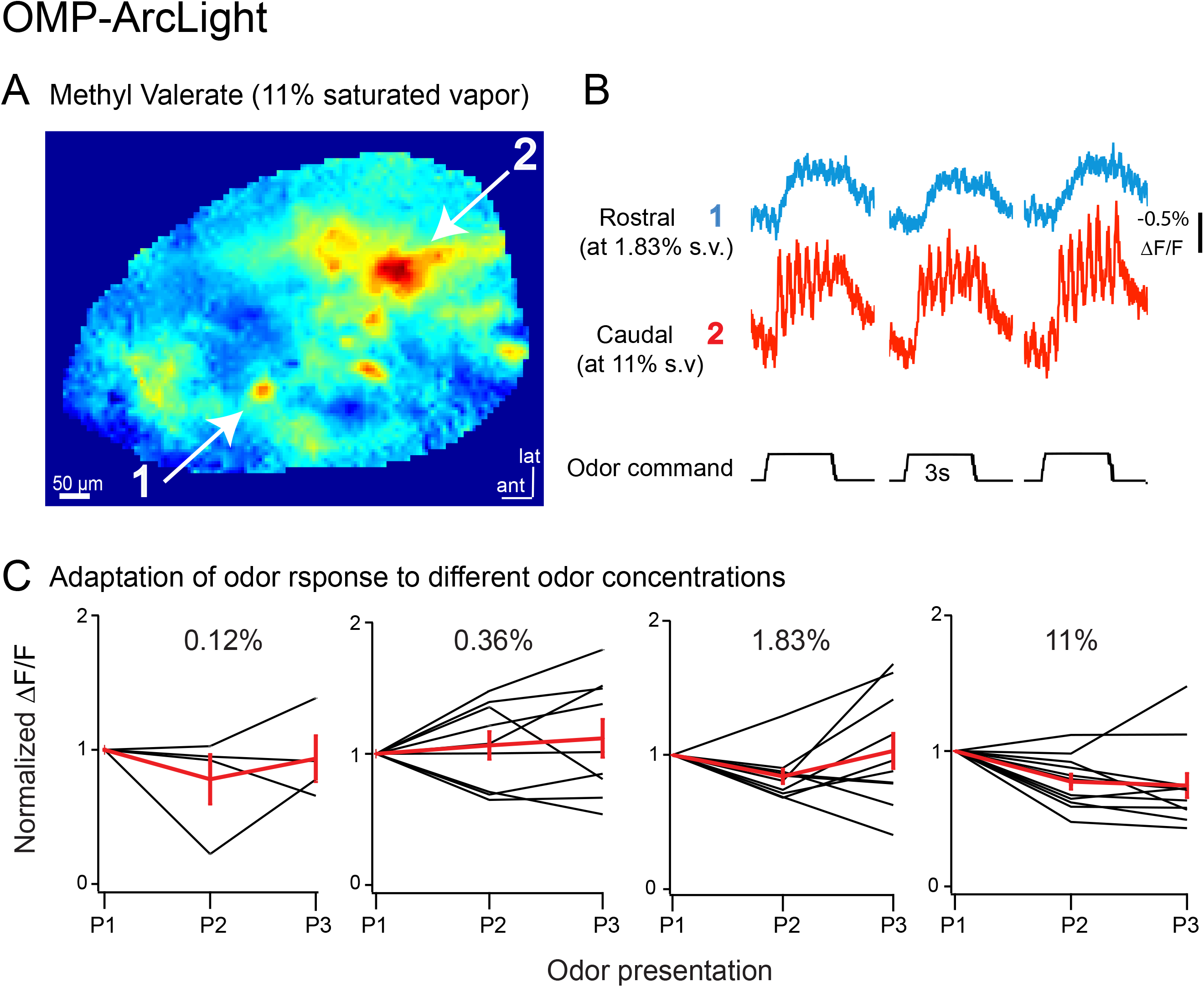
The olfactory bulb input shows little adaptation in response to consecutive odor presentations. **A.** Activity map of neuronal activity in response to 11% of saturated vapor (methyl valerate) with examples of active glomeruli located in rostral [15] and caudal (2, white arrows) part of the olfactory bulb. Scale bar: 50 μm **B.** Optical traces show temporally diverse odor-evoked activity of ArcLight detected in two different glomeruli designated in panel A. The cropped traces of responses to three consecutive odor presentations (3s long) are shown. **C.** A series of graphs showing normalized signal size vs. stimulus presentation[s] at the four different concentrations of methyl valerate. The responses on the second (P2) and third (P3) odor presentation were normalized to responses recorded on the first presentation (P1). The odorant concentration is indicated in the graph. The values from a single preparation are shown in grey. The mean of all responses is shown as a red line, with s.e.m as error bars.

### Emx1-ArcLight mice

Similar imaging experiments using epifluorescence microscopy were carried out using the Emx1-ArcLight transgenic line. Odors evoked patterns of activity that were diffuse and not clearly glomerular (data not shown), a result that was consistent with the expression evident in the histology (Figure 1D). However, different odors (methyl valerate, isoamyl acetate, and ethyl tiglate) resulted in distinctive (but diffuse) activity maps that were relatively invariant with respect to odor concentration (data not shown). Thus, the odor concentration dependence is different from that of the input olfactory receptor neurons (Figure 2) but similar to that of the mitral/tufted output neurons [11]. Furthermore, despite the lack of clear glomerular activity patterns, similar temporal differences were evident in the caudal versus rostral bulb. The temporal differences evident at the OSN level are maintained at the level of the granule cell layer.

#### Two-photon imaging from Emx1-ArcLight mice

Although histological examination confirmed that individual cells were labeled in the glomerular layer in EMX-ArcLight mice, we could not resolve them in epifluorescence imaging due to light scattering and diffuse fluorescence originating from the deeper layers. To address whether voltage imaging could be carried out from these neurons in the glomerular layer, we used 2-photon imaging, which greatly reduces the contribution from signals above and below the focal plane.

Figure 5 illustrates the resting fluorescence intensity at a low magnification (Figure 5A) and with additional magnification (Figure 5B). Glomeruli had relatively weak fluorescence (Figure 5A, #2), although regions surrounding the glomeruli were in part brightly labeled, and appeared as individual neurons with membrane localized fluorescence (Figure 5B, #1). Odors evoked signals that could be detected in single trials (Figure 5C-D) in some of the neurons (e.g., Figure 5B, top, arrow #1). Some signal was also detectable in nearby neuropil (Figure 5D, compare #1 and #3), likely due to GEVIs being expressed all along the membrane of a cell. Similar results were obtained in nine preparations.

**Figure 5.**
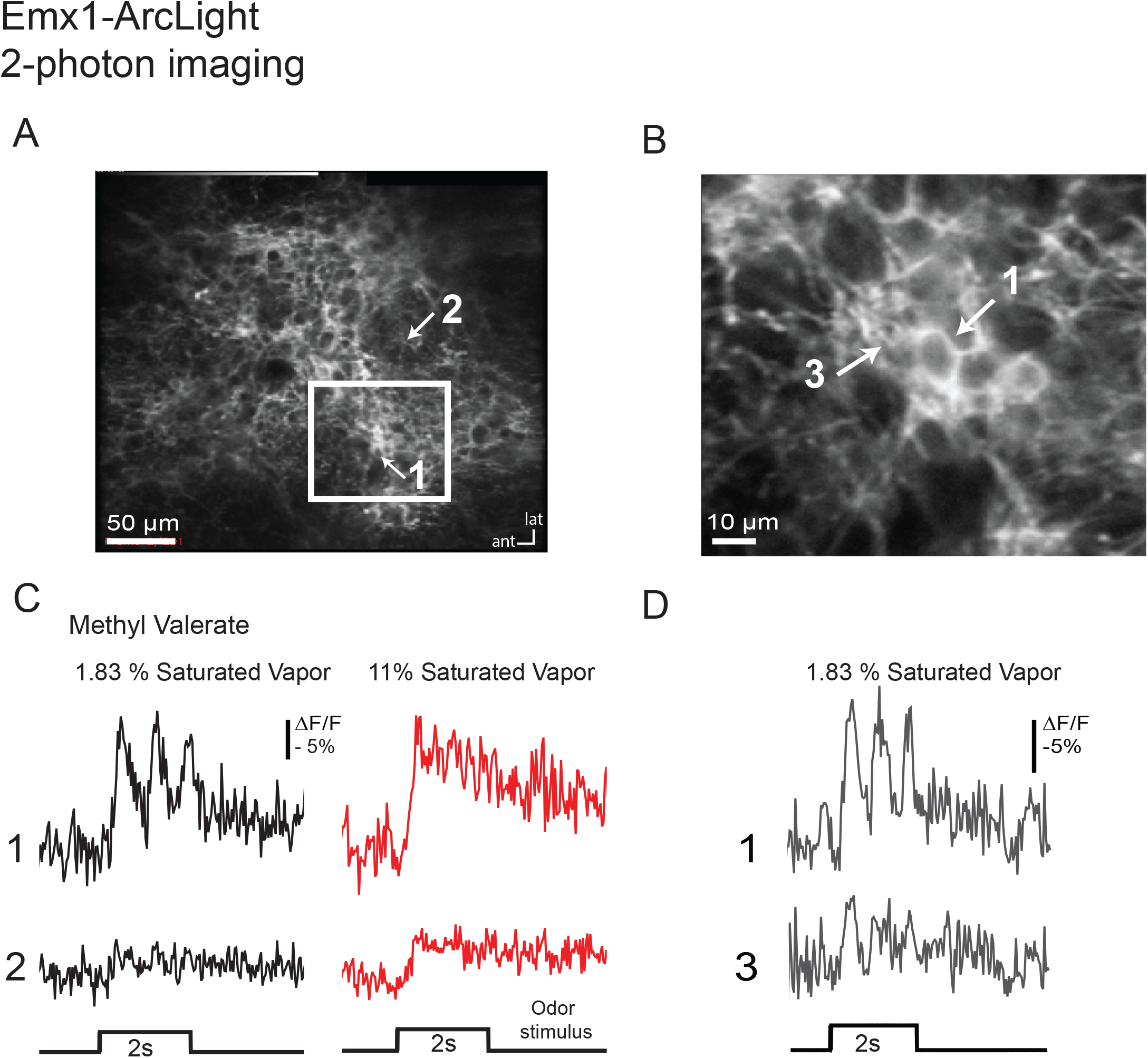
Functional characterization of ArcLight activity in the EMX-ArcLight transgenic mouse line using 2-photon microscopy. **A.** A low magnification resting light image showing membrane-localized expression of ArcLight. **B.** A high magnification resting light image showing membrane-localized expression of ArcLight. **C.** Optical traces showing activity recorded from regions shown by the arrows in panel A. The activity is recorded in response to two different odorant concentrations (2% and 10% of methyl valerate). **D**. Optical traces showing activity recorded from regions shown with arrows in panel B. The signals are unfiltered, single trials.

## Discussion

The GEVI ArcLight is a chimeric voltage indicator [7] made of a voltage sensitive domain derived from the *Ciona intestinalis* voltage sensitive phosphatase [33, 34] and a mutated version [7] of super ecliptic pHluorin, a GFP (A227D) [35, 36]. Neuronal cells expressing ArcLight exhibit a depolarization-dependent decrease in fluorescence intensity. While ArcLight has relatively slow kinetics (~10ms fast time constant), which limits its ability to detect high-frequency spiking activity, it is extremely bright, sensitive, and has been shown to respond to neuronal activity *in vivo*. This includes *in vivo* application in *Drosophila*, worm, mouse, and even plants [10, 11, 17, 18, 37–39]. Previous imaging studies using ArcLight in mice used either *in utero* electroporation or AAV viral vectors under the control of different promoters (hSyn and CAG).

The ArcLight transgenic mouse line, Ai86(TITL-ArcLight; JAX Stock No. 034694), was developed as a part of a larger effort to increase cell-type specific expression of various reporters [21, 23]. Here, we used epifluorescence and 2-photon microscopy to functionally characterize two *de novo* generated mouse lines with distinct patterns of ArcLight expression in the olfactory bulb (OMP- and Emx1-ArcLight). These mouse lines allowed us to examine different aspects of olfactory bulb physiology, such as an odor-evoked activity maps, temporal dynamics of the activity, concentration invariance, and adaptation.

This study reports the first use of voltage imaging to measure odor-evoked responses from mammalian olfactory sensory neurons. Our prior attempts to use nasal infusion of voltage dyes to record from the olfactory sensory neurons failed (unpublished observation, D.A.S. and L.B.C.). Furthermore, attempts to use several AAVs expressing various indicators to transduce olfactory sensory neurons via nasal infusion resulted in little to no expression (unpublished observation, D.A.S. and L.B.C.). Thus, OMP-ArcLight transgenic mice represent the first successful approach to measure membrane potential changes from olfactory sensory neurons in the intact mammalian brain.

Because previous measurements of the odorant concentration dependence and adaptation in the olfactory sensory neurons had been carried out with calcium sensitive indicators [11, 31], there was concern that those results might be dependent on the use of calcium as a surrogate for membrane potential signals. Using the OMP-ArcLight mice we found that the results using the voltage indicator (Figures 2 and 4) were essentially identical to those reported using calcium indicators.

Future experiments will examine voltage imaging targeted to other cell types, which will allow a more temporally precise characterization of the input-output transformation of the olfactory bulb.

## Acknowledgements

We thank Yunsook Choi for helpful comments on the manuscript. We would also like to thank The John B. Pierce Laboratory, Inc., Florida State University, and KIST for ongoing support. We would like to acknowledge the expert technical contributions of the Pierce Laboratory Instrument shop including John Buckley, Andrew Wilkins, Tom D’ Alessandro, Ronald Goodman and Angelo DiRubba. The authors have no competing interests to report. This study was supported by NIH grants U01 NS090565, R01 NS083875, R01 DC005259, U01NS103517 and UF1NS107705.

